# Characterization of neural communication dynamics in the Ventral Attention Network across distinct spatial and spatio-temporal scales

**DOI:** 10.1101/2019.12.25.888446

**Authors:** Priyanka Ghosh, Dipanjan Roy, Arpan Banerjee

## Abstract

The Ventral Attention Network (VAN) is involved in reorienting attention from an ongoing task when a salient (pop-out) stimulus is detected in the environment. Previous neuroimaging studies have extensively evaluated the structural and functional connectivity of the VAN. However, directed effective connectivity within the network and the neural oscillations driving it still remain elusive. Functional magnetic resonance imaging (fMRI) studies have not been able to address this issue due to lack of appropriate temporal resolution required to capture the process of reorientation. In this study, we recorded scalp electroencephalography (EEG) and behavioural data from healthy human volunteers, obtained saliency-specific spectral changes, localized the sources underlying the spectral power modulations with individual-specific structural MRI scans, reconstructed the waveforms of the sources and investigated the causal relationships between the areas of the VAN using Granger causality (GC). Using a custom-designed experiment involving visual search on static images and a dynamic motion tracking task, we investigated the neural processing of salient distractors operating at very slow and very fast time scales, respectively. Our results revealed how a task-independent but context-specific VAN encompassing the right insula, the right lateral pre-frontal cortex, the anterior and the posterior right temporo-parietal junction communicating in the alpha frequency band (8-12 Hz) supports saliency processing.

## 1. Introduction

The ability to reorient our attention towards unexpected salient changes around us while performing routine tasks (e.g., spotting a speeding car while crossing the street) is critical for our survival. Where in one hand, goal-directed ‘top-down attention’ is required to orient attention towards a target to achieve a pre-decided goal in a task, response to a ‘pop-out’ or salient stimulus involves the rapid capture of attention by shifting of the attentional focus from an ongoing goal-directed task through what is known as ‘bottom-up attention’ guided by the Ventral Attention Network (VAN) (Corbetta & Shulman, 2002; Corbetta, Patel, & Shulman, 2008). This process of attentional reorientation is essential but transient at the same time as attention needs to be oriented back to the task at hand. Since the brain is limited by attentional resources, synchronization or desynchronization of alpha oscillations (8–12 Hz) might be one way how the brain gates attentional selection on the neural level. One view is that synchronized alpha-band oscillations inhibit and desynchronized alpha-band activity excites sensory cortical areas (or at least reflects these processes) (Klimesch et al. 2007; Jensen and Mazaheri 2010; Foxe and Snyder 2011). Regardless of a vast body of existing literature that has linked alpha oscillations to distractor suppression, the neural implementation of this mechanism of inhibition at the level of attentional networks lacks clarity.

Extant literature implicates the VAN comprising of the anterior insula, the right temporo-parietal junction (rTPJ) and the lateral prefrontal cortical areas (lPFC) comprising inferior frontal/middle frontal gyri (IFG/MFG) to be responsible for processing salient stimuli or oddballs (Allan et al., 2020; Vossel, Geng, & Fink, 2014; Han & Marois, 2014; Corbetta, Patel, & Shulman, 2008). Even though the VAN has been studied in much structural and functional detail, the existing studies offer very limited insight into the directed connectivity within this network. Abnormal VAN function has been associated with many clinical conditions like depression (Liu et al., 2019), which makes the study of this network and the neural processes associated with it important. Till date, only a few fMRI studies have tried to investigate the causal architecture underlying the VAN using dynamic causal modelling (DCM) (Liu et al., 2019; DiQuattro et al., 2014; Vossel et al., 2012). The sampling rate (repetition time or TR) for fMRI is however, very low (in the order of seconds), which is much slower than the temporal resolution (in the order of milliseconds) required to capture the neural responses of such instantaneous attentional shifts. Hence, the study of network dynamics underlying the process of reorientation of attention may be limited by the sluggishness of the BOLD haemodynamic responses. To circumvent this problem, we used high-density EEG with simultaneous behavioural recordings from healthy human participants. Accurate source reconstruction techniques that involved co-registration of EEG with individual subject’s brain MR images were employed to locate the brain networks underlying alpha modulation, followed by characterization of the causal architecture of the constituent regions using Granger causality analysis (GCA). Data obtained from EEG recordings of continuous neural activity are well suited to GCA by virtue of having high temporal resolution.

As evident, attention is just not limited by resources but also by time (Nobre and Coull, 2012). Previous studies have reported that the minimum dwell time for attention at a fixed location is about 200 ms whereas when the focus of attention is changing along with time, a given location on the moving target’s path can be selected for extremely brief time periods approximating to 50 ms (P Cavanagh, Battelli, & Holcombe, 2014). Hence, the study of the neurobiology of attention requires a detailed understanding of the resource-wise allocation of attention both in space and time, depending on the task at hand. For instance, how the brain processes information to execute a visual search over a static image is entirely different from the sensory and cognitive processing deployed in executing a task involving tracking of a dynamic stimulus (Kulikowski & Tolhurst, 1973; Battelli et al., 2001; Battelli, Pascual-Leone, & Cavanagh, 2007; Stigliani, Jeska, & Grill-Spector, 2017). Unlike a static stimulus, attention to a dynamic stimulus has limits extending over space and time, because when the speed of the stimulus increases, tracking ability decreases (P Cavanagh & Alvarez, 2005). Thus, a broader question arises, is the processing of reorientation of attention towards inclement bottom-up signals driven by different or common brain circuits across tasks? We hypothesize that the underlying networks responsible for bottom-up attention are independent of the nature of the task and that such phenomena are also reflected in the modulation of alpha oscillations while processing salient distractors.

To test this hypothesis, we designed two visual attention tasks involving static and dynamic stimulus processing, where the participants were presented with a salient distractor in the same visual space as the target during an ongoing goal-directed task. Such an experimental design was chosen to keep the visual stimulation close to what one would experience in a real life situation. Some previous EEG/MEG studies on bottom-up or exogenous attention are severely limited by their presentation method where the salient distractors were either part of the visual display right from the onset of a trial along with the target or there were separate trials (involving valid/invalid cues) for endogenous and exogenous attention (Carretié et al., 2017; Dugué et al., 2018; Feng et al., 2017; Kiss et al., 2012; Peelen et al., 2004) that were evaluated independently of each other which we believe is an unwarranted measure to study the process of reorientation. The aim of the present study is to determine a direct relationship between neural networks underlying the alpha power enhancement and salient distractors across two completely different task conditions: one involving spatial complexity and the other spatio-temporal complexity, on a comparative scale. Our study also seeks to reveal the causal relationships between the various regions of the VAN involved in saliency processing through a time-varying Granger causality approach, again comparing it across tasks spanning spatial and spatio-temporal scales.

## 2. Methods

### 2.1. Participants

22 healthy human volunteers (11 females and 11 males) aged between 21-29 (mean = 26.9, SD = ±2.15) years were recruited for the study. All participants had University degrees or higher; were right handed (indexed by laterality score according to the Edinburgh handedness questionnaire); reported normal or corrected-to-normal vision; and declared no history of neurological or psychiatric disorders. The participants were requested to avoid the intake of any stimulant or medication (e.g., coffee, sedatives etc.) before coming for the experiment. The study was carried out following the ethical guidelines and prior approval of the Institutional Review Board of National Brain Research Centre, India which conforms to the standard set by the Declaration of Helsinki. Written informed consent was obtained from all participants before the commencement of the experiment and they were remunerated for the time of their participation.

### 2.2. Rest block

Before starting with the experimental task, five minutes of eyes open resting-state EEG data were collected from the participants. During this period, a blank black screen was presented on the monitor. The participants were asked to relax or think at free will while viewing the monitor screen placed before them. They were requested to make minimal head, body and eye movements.

### 2.3. Stimulation blocks

All the participants performed two visual attention-based tasks which incorporated two stimulus conditions: static and dynamic (Figure 1). The entire experiment was divided into 16 blocks (8 blocks of each stimulus condition). Each block was presented in a random order during the experiment but was never repeated. Both the visual tasks had three categories of trials: ‘Without Saliency Trials’ **(WT)**, ‘Saliency Trials’ (**ST**) and ‘Neutral Trials’ (**NT**). The presentation order of the three categories was randomized in each block. The participants were not aware of the categorization in trials and were instructed only about the respective goals of the static and dynamic tasks before the experiment.

**Figure 1.**
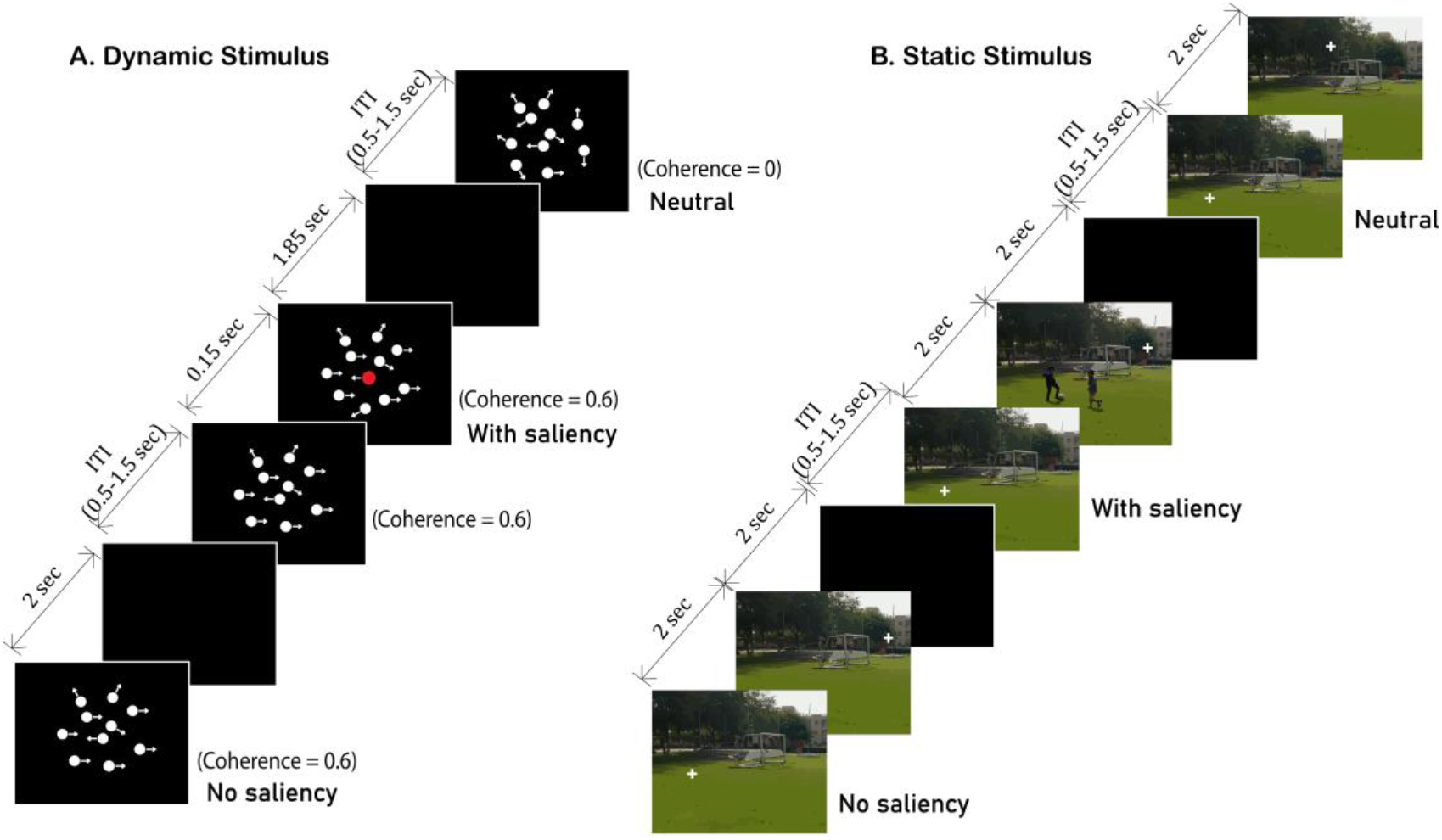
Experimental design. An example of the experimental paradigm designed for the study illustrating the three different categories of trials: neutral trials (NT), without saliency trials (WT) and saliency trials (ST) is shown along with their presentation durations within a block which comprised of videos in the (A) Dynamic stimulus condition and static images in the (B) Static stimulus condition.

The NT served as a control to the participants’ attention. They were introduced to keep a check if the participants were attentive throughout the experiment and were just not making random responses. The NT were designed to give an impression of the most difficult trials to the participants, which if attended, were expected to produce the longest reaction times. The distribution of trials within a block for both the static and dynamic visual tasks is given in Table 1.

**Table 1.** Trial distribution across tasks. The block-wise distribution of the Neutral Trials (NT), Without Saliency Trials (WT) and Saliency Trials (ST) across the dynamic and the static stimulus conditions have been listed.

| **Trial information** | **Dynamic stimulus** | **Static stimulus** |
| --- | --- | --- |
| *Total no. of blocks* | 8 | 8 |
| *No. of trials per block* | 70 | 30 |
| *NT* | 20 | 10 |
| *WT* | 20 | 10 |
| *ST* | 30* | 10 |

\*To reduce the drop in the pop-out effect of salient distractors due to habituation after multiple trial presentations, 3 kinds of salient distractors were used, varying in either color or size or both from the other moving dots. 10 trials each of an equisized red, a larger red and a larger white dot were presented in a block as a salient distractor along with rest of the moving dots in ST.

Inter-stimulus intervals (ISI) between successive trials were randomly drawn from a uniform distribution with values ranging between 500 ms and 1500 ms (mean = 1000ms) in which a blank black screen was presented to avoid any saliency-related effects due to central fixation.

Stimulus presentation and behavioral response collection were done using Neurobehavioral Systems (NBS) Presentation software. Participants viewed the stimuli on a 21’’ LED screen (1280 × 1024 pixels) with a 60 Hz refresh rate placed on a 74-cm-high desktop. The center of the screen was placed within 10–20° of the participant’s line of sight, at a 60–70 cm distance. The stimuli were presented on a black background over which the static stimulus covered an area of 20 × 20 cm on the screen whereas the diameter of the aperture in the dynamic stimulus was 20 cm.

#### Dynamic stimulus

The dynamic stimulus viewing task was a four-alternative forced choice (4-AFC) task. The stimuli were designed using Psychtoolbox-3 in MATLAB R2016b and were exported as videos with a frame rate of 60 Hz. The participants were presented these videos which consisted of white-colored equal-sized randomly moving dots where, a proportion of dots moved in a particular direction because of a certain coherence assigned to them. The coherence of the dots was kept at 0.6 for all the trials, which means that out of 100 dots, 60 dots moved in one specific direction and the other 40 moved in random directions, uniformly distributed over 0-360 degrees. The speed of motion of all the dots was kept constant across all trials. The participants were instructed to identify the net direction of the moving dots which could either be left/right/up/down and respond using the respective arrow keys on the keyboard. Each video was presented for 2000 ms. The goal in the task was the same for both WT and ST, with the only difference in the latter being the emergence of a salient dot at a timestamp of 150 ms from the onset of the trial, moving randomly within the same aperture as the other dots. The 150 ms latency was decided with the purpose of creating an interference in the decision-making process (Teichert, Grinband, & Ferrera, 2016) of the participant while doing the goal directed task. The experimental schematic is illustrated in Figure 1.

#### Static stimulus

The static stimulus consisted of a two-alternative forced choice (2-AFC) task. The participants were presented with two similar pictures on the screen, successively. Each picture pair made up one trial and was randomly selected from a pool of twenty such picture pairs. Thirty such picture pairs were presented in one block. The pictures were naturalistic images (from both indoor and outdoor settings; no faces included) captured using a 16 MP camera keeping the settings same for all images. Using Adobe Photoshop CC 2015.5, a white-colored ‘+’ shape (size 1/800th of the image) was added to all the images at random positions. Multiple copies of a single image with ‘+’ shape at different positions were created such that there was no image and ‘+’ position memory association. Each picture was presented for 2000 ms. This was a visual search task where the participants had to search for the white-colored ‘+’ shape in both the pictures and report its change in position in the second picture with respect to the first picture. For convenience, the participants were advised to imagine a vertical line bisecting the screen into left and right halves. They were instructed to press the upward arrow key if the ‘+’ sign moved to the same half of the screen in the second picture, i.e, change in position was on the same side of the imaginary line; and to press the downward arrow key if the ‘+’ sign changed its position and moved to the other half of the screen i.e, from the left half to the right half or vice versa. The goal in the task was same for WT and ST, with the only difference in the latter being the introduction of a salient (‘pop-out’) object in the second picture at any random position. An example of one such stimulus is presented in Figure 1.

### 2.4. EEG Data Acquisition

Behavioral and EEG data were acquired in the EEG recording room where ambient noise, light, and other interferences were strictly controlled during the experiment to the same levels for all recording sessions. A Neuroscan EEG recording and acquisition system (Scan 4.3.3 & Presentation), which included an elastic cap (EasyCap) with 64 Ag/AgCl sintered electrodes and amplifier (SynAmps2), was used. The 64-channel EEG signals were recorded according to the International 10–20 system of electrode placement. Cz was the reference electrode, grounded to AFz and the impedance of all channels was kept below 10 kΩ. The data were acquired at a sampling rate of 1000 Hz. A Polhemus Fastrak system was used to record the 3D location of electrodes using a set of fiducial points (Cz, nasion, inion, left and right pre-auricular points) while the EEG cap was placed on the participant’s head.

### 2.5. Behavioral Data Acquisition

All the responses were made on a computer keyboard using left/right/up/down arrow keys and were recorded through the NBS Presentation software by receiving triggers at keyboard presses. Before the experiment, the participants were instructed to watch the stimulus carefully before making any response. They were asked to be as fast and as accurate as possible and respond to an ongoing trial before it’s offset. A blank screen followed by the subsequent trial appeared automatically after the offset of the ongoing trial, regardless of whether the participants had responded or not. They were also asked to respond to all the trials. If more than one response was made for a trial, only the first response was considered for further analysis. A rest period was allowed after every block, with the participant deciding the length of the rest period to maintain minimum fatigue.

### 2.6. EEG Data Preprocessing

For both the static and the dynamic tasks, pre-processing steps and analysis pipeline were identical. All the pre-processing steps were done with the EEGLAB toolbox (Delorme & Makeig, 2004) and custom-written scripts in MATLAB (www.mathworks.com).

Raw EEG data from all the participants were imported using EEGLAB toolbox (Delorme & Makeig, 2004) following which they were first filtered using a band-pass filter of 0.1-80 Hz followed by a notch filter between 45-55 Hz to eliminate line noise at 50 Hz. Post filtering, the data were visually inspected and the trials with any abnormal or noisy segments (jitters with very large amplitudes) were removed. Data of two participants were discarded at this step due to very noisy recordings. Next, the filtered data were re-referenced to the common average. Epochs of 1000 ms post salient stimulus onset were extracted using trigger information and were sorted from WT, ST and NT categories. Trial-by-trial detrending of each epoch category was performed to remove linear trends from the signal. To further remove ocular, muscular and electrocardiograph artifacts, a threshold of ±75µV was set and trials with a magnitude beyond this threshold at any time-point were rejected from all the channels. Overall, about 70% of the trials for each task condition from each subject were preserved after artifact rejection.

### 2.7. Behavioral Analysis

The reaction times and accuracies of all the trials were calculated. Since attention is a key component in our experiment and any form of distraction (internal/external) could shift attention away from the task, blocks with response accuracies less than 70% (less than 6% of all blocks) were excluded from further analysis. To decrease the number of false positives and minimizing chances of including responses made without the involvement of attention, all incorrect trials including the skipped trials were also excluded. Data from one participant were discarded due to very poor performance specifically in dynamic task trials. To rule out the possibility of incorrect responses being made because of a specific directional bias in any participant, we computed the percentage of incorrect responses for each direction and found that it was nearly the same for all directions in each individual.

From the remaining 19 participants, the reaction times of the correct trials were sorted and averaged across all the participants for both static and dynamic tasks. The reaction time was the duration from the onset of the stimulus till the participant hit the response button. For the NT, any response was considered as correct.

### 2.8. Spectral Analysis

To understand the neural correlates of the behavior and hence, the processing of saliency by the brain, we looked at the constituent frequencies from 0.1-80 Hz in the individual trial categories of both the tasks. Power spectrum analysis was performed on the EEG time series data for the 19 participants. To ensure an equal contribution of trials from each participant, the number of trials from the participant with the minimum number of remaining trials after artifact rejection was chosen for further analysis. Those many trials were randomly sampled and extracted from all the three categories for each participant. Since 35 was the minimum number, we had a total of 665 (19*35) trials from each category in both static and dynamic tasks.

The analysis scheme was designed in a way that could tease out the effect of the salient distractor while doing the goal-directed task and therefore, a time window of 1000 ms from the onset of saliency was considered (timestamps were matched accordingly for WT and NT). Using the EEG time-series data, we calculated the power spectral density for each trial corresponding to all the 64 channels using the multi-taper method (mtspectrumc.m) provided by Chronux toolbox (Bokil et al., 2010). A standard multi-taper FFT was used that applied 5 Slepian tapers to each window (time bandwidth product = 3). Sampling frequency was kept at 1000 Hz and frequencies were estimated between 0.1 to 80 Hz.

### 2.9. Source Reconstruction using individual T1 MRI images

To localize the sources of the alpha band activity, we applied a current density technique: exact low-resolution brain electromagnetic tomography (eLORETA) implemented by the MATLAB-based Fieldtrip toolbox. eLORETA (Pascual-Marqui, 2007) is a weighted minimum norm inverse solution that provides exact localization with zero error in the presence of measurement and structured biological noise. We first created the forward models of individual participants using their respective T1-weighted structural MRI images (MPRAGE) collected from a Philips Achieva 3.0 T MRI scanner using the following acquisition parameters: TR = 8.4 ms, FOV = 250 × 230 × 170, flip angle = 8 degrees, and fiducials marked at nasion, left and right pre-auricular points with Vitamin E capsules. The origin of all the T1 images was set to the anterior commissure using SPM 8 before generating individual head models. Using Boundary Element Method (BEM), the brain was segmented into a mesh/grid based on the geometrical and tissue properties of the brain. The Polhemus data with the electrode locations of individual subjects, was then fitted over these individual head models co-registered to the MRI fiducial points to create the leadfield matrix corresponding to each participant. For a frequency-domain source analysis, the cross-spectral density (CSD) matrix, which contains the cross-spectral densities for all sensor combinations, was computed for individual participants from the Fourier transformed data for the alpha frequency band (8-12 Hz).

Using the CSD matrix and the lead field matrix, a spatial filter was calculated for each grid point. By applying this spatial filter to both the trial conditions (WT and ST) separately, the power estimate for each grid point was obtained. For calculating the source power, a common filter approach was used to ensure that the differences in source power across the two trial conditions were actually because of differences in the brain activity and not because of differences in the filter output (which might arise due to variations in the signal-to-noise ratio and subsequently varying CSD matrices) in the two trial conditions (WT and ST). Using this common inverse filter, the net source power was computed for each participant and the individual grids were interpolated with their respective T1 weighted images followed by normalization over a common Colin 27 brain template. The statistical threshold was set at 99% significance level to define activated sources of alpha enhancement.

### 2.10. Source Time-series Reconstruction

From the thresholded grid points, we reconstructed the time series for the activated ROIs at the source level by multiplying the spatial filter with the artifact rejected time-series data using common electrode placements for all individuals computed by taking the average of their normalized 3-D locations. The projection of the filter onto the EEG time series data for each task condition yielded 3 source dipole time-series with their orientations along the x, y and z directions. Since the interpretation of results becomes difficult while dealing with three dipole orientations, the time-series were projected along the strongest dipole direction. This was done by determining the largest (temporal) eigenvector corresponding to the first singular value. Using these reconstructed time series, connectivity analyses were done for each task condition to look for effective connectivity between the sources involved in processing saliency.

### 2.11. Effective Connectivity Analysis

To understand the directional interactions between the sources, effective connectivity was calculated using conditional Granger Causality in the time domain using a Causality Estimating Software (https://www.dcs.warwick.ac.uk/~feng/causality.html). This software uses a time-varying Granger Causality approach catered to deal with non-stationary time series of EEG/MEG data, unlike the traditional Granger Causality methods that fit a time-invariant multivariate autoregressive model (MVAR) to the time series.

The reconstructed time-series from each participant was treated as a trial, leading to reconstruction of 19 trials from 19 participants. One participant’s data from the dynamic stimulus was removed as the reconstructed time series data was very noisy. To further reduce non-stationarity, all the reconstructed time series were bandpass filtered between 5-45Hz to minimize the effects of evoked potentials. A connectivity matrix was created between all the nodes (7 × 7 for dynamic stimulus and 5 × 5 for static stimulus). The time series data for each node was subjected to 50 rounds of bootstrapping from which the mean Granger causality value for each causation combination was obtained. To test the statistical significance of these values, a 95% confidence interval was generated empirically from a null distribution by random permutation of the time series across all the nodes and trials for 50 times. Subsequently, Granger causality estimates were obtained for these 50 iterations, and the mean and the standard deviation were computed from the GC values to compute the confidence interval at 95% significance.

### 2.12. Data availability & Codes

Raw data and processed data used to generate the figures will be made available by the corresponding author on request.

## 3. Results

### 3.1. Behavioral performance

The mean and standard error of the mean of the reaction times of all the trial categories NT, WT and ST are shown in Figure 2 for dynamic and static tasks. Statistical significance was computed using Wilcoxon rank-sum test where the null hypothesis of no significant difference between reaction times of any two categories in a task condition was tested at 95% confidence level. The reaction times showed a significant difference (NT>WT at p<0.0001, NT>ST at p<0.0001 and ST>WT at p=0.012 in dynamic task; NT>WT at p<0.0001, NT>ST at p<0.0001 and ST>WT at p<0.0001 in static task) between each trial category within a task and followed a similar pattern of reaction time relationship in both the tasks which was: NT >ST > WT.

**Figure 2.**
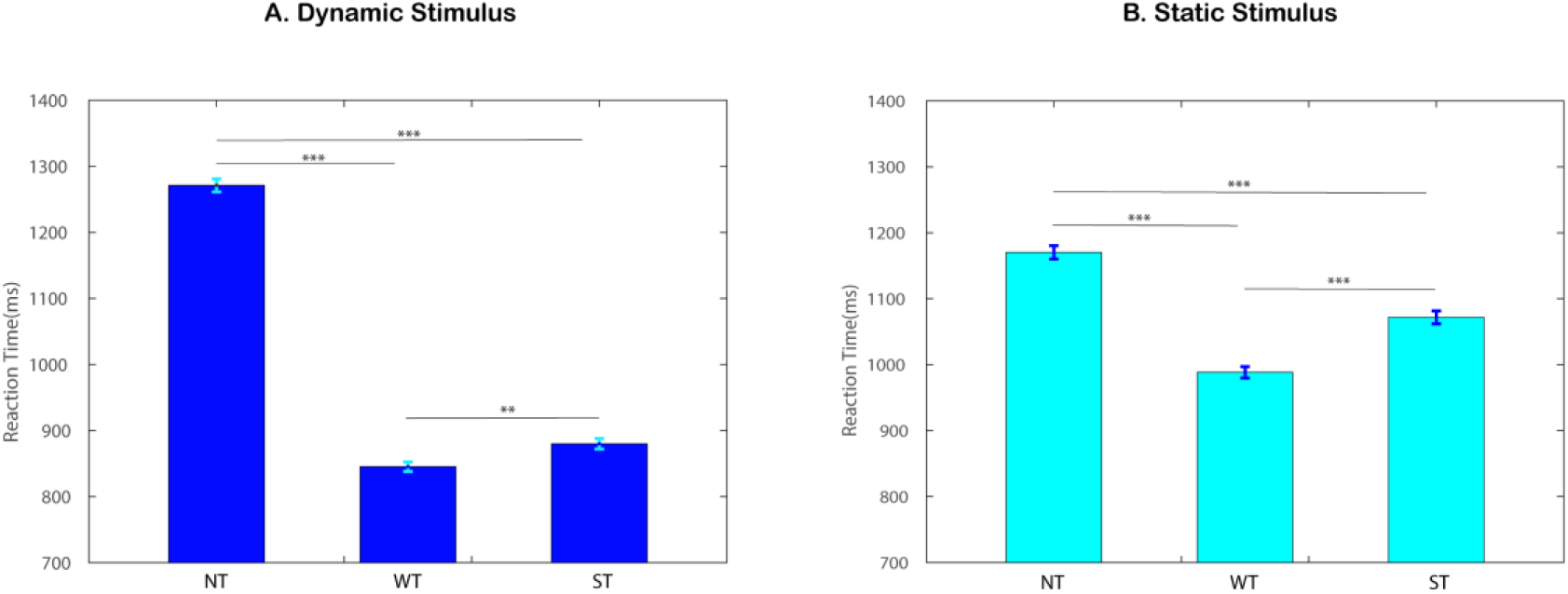
Behavior. The mean and standard error of the mean of neutral trials (NT), without saliency trials (WT) and saliency trials (ST) from all the nineteen participants are shown for the two task conditions: (A) Dynamic stimulus condition and (B) Static stimulus condition. The significant difference between any two categories of trials within a task condition was tested at 95% confidence interval using Wilcoxon ranksum test (*** represents p<0.0001 and ** represents p=0.012).

### 3.2. Spectral power estimates

The power spectra were calculated for WT, ST and NT of all the participants trial-by-trial, which was subsequently averaged and collapsed across all the 64 sensors for both the task conditions: dynamic and static. Figure 3 compares the normalized power spectra for ST, WT and NT. Though all the trial categories in a task followed the same pattern of power spectra, the power of the alpha frequency band (8-12 Hz) was higher for the ST as compared to the WT and the NT. Interestingly, this pattern was seen in case of both dynamic and static stimuli. To test out the statistical significance of the increased alpha power, we employed a non-parametric Wilcoxon rank-sum test which revealed that alpha power significantly increased in ST as compared to WT (p=0.002) and NT (p<0.001) for the static stimulus condition between 8 to 11 Hz at 95% confidence level. Similarly, in the dynamic stimulus condition, there was a significant increase (p = 0.04) in the alpha power of ST as compared to WT between 8 to 11 Hz. The increase of power in ST as compared to NT was however, marginally significant (p = 0.06). To verify the robustness of this alpha enhancement pattern and find the peak alpha frequency, we employed a detrending method and compared the alpha power spectra of NT, WT and ST conditions. Here, we modeled the 1/f trend of the log-transformed power spectrum and subtracted this trend from our original power spectrum data. This detrended power spectrum was then plotted for the entire alpha frequency range between 7 to 14 Hz. Interestingly, the enhanced power of ST had the peak frequency at 10 Hz for both the static and the dynamic tasks (**insets to** Figure 3).

**Figure 3.**
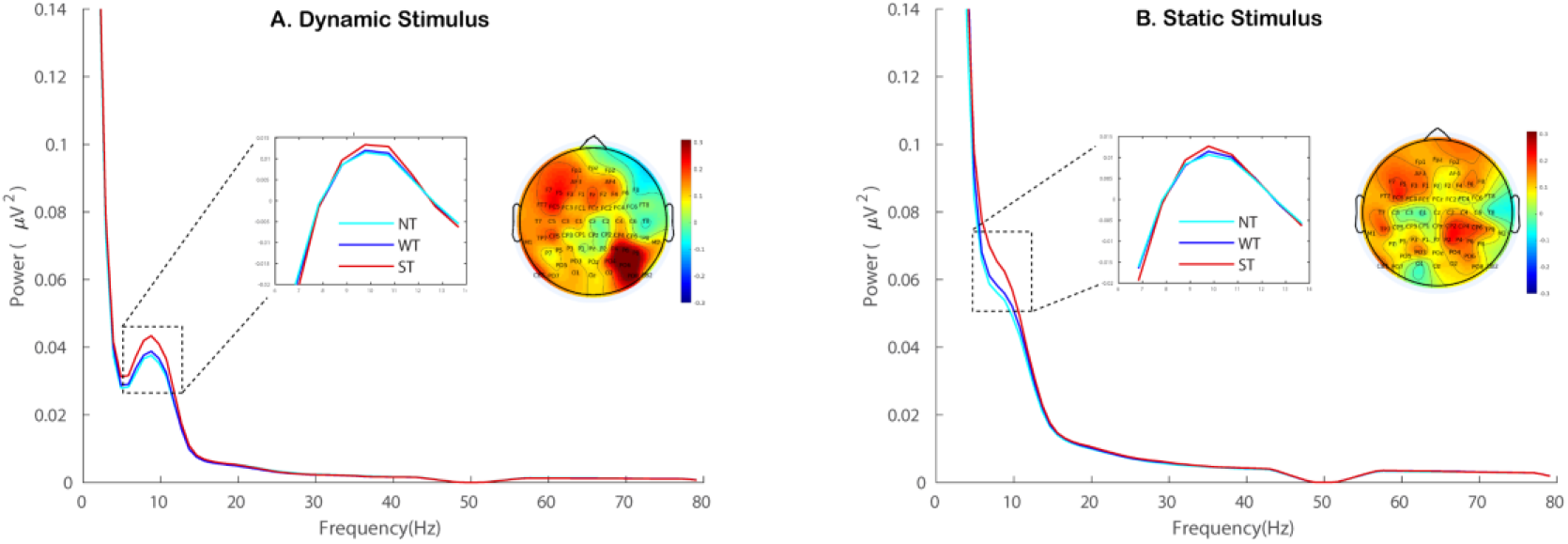
Power spectral density. The figure shows the grand-average of the power spectra of neutral trials (NT), without saliency trials (WT) and saliency trials (ST) across (A) Dynamic Stimulus condition and (B) Static Stimulus condition. The boxed regions represent those frequencies (alpha band, 8-11 Hz) which show a significant increase of power (ST>WT at p=0.002 and ST>NT at p<0.0001 in static stimulus condition; ST>WT at p=0.04 and ST>NT at p=0.06 in dynamic stimulus condition) in ST as compared to WT and NT, validated using Wilcoxon rank-sum test for individual frequency pairs within a condition. The left inset in each stimulus condition shows that the peak of alpha is at ∼10 Hz for both (A) Dynamic Stimulus and (B) Static Stimulus. The right inset in each stimulus condition are topoplots highlighting the enhancement in alpha peak power (∼10Hz) in ST with respect to WT across the sensor space computed using the alpha modulation index (AMI).

The topoplots for dynamic and static stimulus processing conditions were plotted (**insets to** Figure 3) for the peak alpha frequency (∼10 Hz) using the formula for alpha modulation index (AMI) (Sokoliuk et al., 2019).

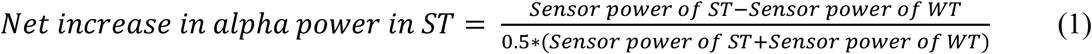

Sensors Fz, F3, F5, F7, FC5, FT7, CP5, TP7, CP4, CP6, P4, P6, P8, PO4, PO6, PO8 and O2 showed the maximum increase in their peak alpha powers at 10 Hz in ST in the dynamic task. Similarly, sensors AF3, F1, F3, F5, F6, F7, FC5, FT7, T7, TP7, CP2, CP4, CP6, TP8, P2, P4, P6, PO3, PO4 and PO6 showed the maximum alpha power increase at 10 Hz in the static task. Overall, in both the tasks, enhanced alpha power concentrated around the centro-parietal, parietal, parieto-occipital and temporo-parietal sensors on the right; and on the frontal, fronto-central and temporo-parietal sensors on the left.

### 3.3. Underlying cortical sources of neural activity

The underlying sources responsible for the enhanced alpha power in ST with respect to WT were calculated using the same formula of alpha modulation index (AMI) at the peak alpha frequency (∼10 Hz) after computing the individual sources for ST and WT using eLORETA (as described in methods).

The relative difference in source powers during ST and WT conditions produced the residual source powers. We argue that the dynamic or static task-specific information was thus negated and the residuals reflect the effect of saliency only. The source powers for all participants were grand-averaged and tested for statistical significance. The grid points that survived 99^th^ percentile threshold, were considered as significant sources of activation in response to salient distractors. For the purpose of plotting, the source coordinates in the 3-D voxel space were projected to a surface plot as represented in Figure 4 using customized MATLAB codes. Spurious activations towards the center of the brain arising from noise were removed by masking the grid points deep inside the brain with an ellipsoid of optimum radii centered at the anterior commissure.

**Figure 4.**
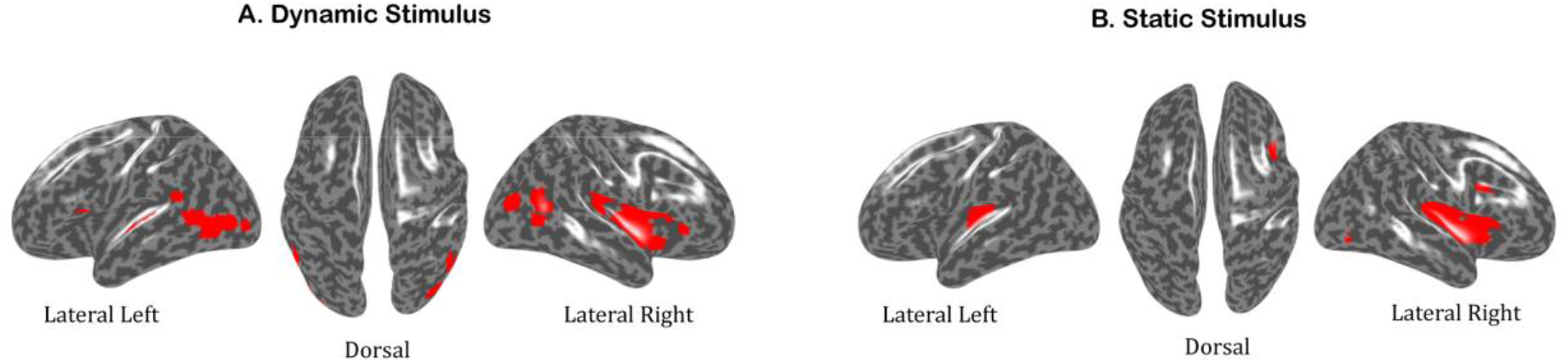
Source localization. The figure represents the cortical sources responsible for processing a salient distractor at the peak alpha frequency (∼10 Hz) computed using the alpha modulation index (AMI). The sources that were identified using eLORETA, a weighted minimum norm inverse solution, were the left and the right anterior temporo-parietal junction, the right posterior temporo-parietal junction, the right insula, the right lateral prefrontal cortex, the left and the right visual association areas for (A) Dynamic Stimulus; and the left and the right anterior temporo-parietal junction, the right insula, the right lateral prefrontal cortex (including the inferior frontal gyrus), the right visual association areas for (B) Static Stimulus. All the regions were approximated to the nearest Brodmann areas of the human brain.

The underlying sources of alpha enhancement in the dynamic task were the left and the right anterior temporo-parietal junction (supramarginal gyrus), right posterior temporo-parietal junction (angular gyrus), the right insula, the lateral prefrontal cortex and regions from the left and the right visual cortex.

The sources corresponding to alpha power enhancement in static task were the left and the right anterior temporo-parietal junction (supramarginal gyrus), the right insula, the right lateral prefrontal cortex (including the inferior frontal gyrus) and regions from the right visual cortex.

Since the reconstructed time series obtained using the thresholded sources consisted of very few (∼1000) grid points from the entire brain, we reconstructed the sources again with a lower threshold (98^th^ percentile), to get more number of grid points. Even though the number of grid points increased upon lowering the threshold, the anatomical landmarks corresponding to source locations did not show much difference overall. Further, using k-means clustering, these sources were classified into nodes based on the centroid of the sources. Sources corresponding to the dynamic task were classified into 7 nodes whereas for the static task, they were classified into 5 nodes. All the regions corresponding to these nodes with their respective coordinates (approximated to the nearest Brodmann areas) have been listed in **Table 2**.

**Table 2.** Areas involved in processing saliency. Coordinates of the regions of interests (ROIs) of the reconstructed sources involved in saliency processing for a) dynamic stimulus condition and b) static stimulus condition are listed.

| **MNI coordinates of ROIs of dynamic stimulus** |  |  |  |  |
| --- | --- | --- | --- | --- |
| *x* | *y* | *z* | *Brodmann area* | *ROI name* |
| 45.5304 | 1.8176 | 7.6352 | 45 | R. LPFC/IFG |
| 49.0328 | -67.0187 | 6.234 | 19 | R. visual area |
| -43.0262 | -44.5205 | 8.2927 | 40 | L. ant. TPJ / SMG |
| -52.8281 | -71.4317 | 0.9335 | 19 | L. visual area |
| 43.7931 | -56.3395 | 28.768 | 39 | R. post. TPJ / AG |
| 47.7159 | -14.6687 | -0.9942 | 13 | R. insula |
| 47.8512 | -30.7366 | 16.2046 | 40 | R. ant. TPJ / SMG |

b).
| **MNI coordinates of ROIs of static stimulus** |  |  |  |  |
| --- | --- | --- | --- | --- |
| *x* | *y* | *z* | *Brodmann area* | *ROI name* |
| 44.6437 | -17.0274 | 13.1784 | 40 | R. ant. TPJ / SMG |
| -40.6605 | -32.5528 | 11.1861 | 40 | L. ant. TPJ / SMG |
| 45.2284 | 0.0653 | -2.7002 | 13 | R. insula |
| 41.0606 | 15.3615 | 18.6218 | 9 | R. LPFC |
| 41.8174 | -75.7257 | -3.1578 | 19 | R. visual area |

The observed alpha power enhancement during dynamic stimulus processing correspond to two sub-regions of the right TPJ: the anterior and the posterior right TPJ. To further confirm those were not just two clusters obtained from a single big region due to a limitation of the clustering algorithm, we calculated the Euclidean distance between the right anterior and right posterior TPJ based on their coordinates, which was equal to 28.80 mm, which showed that the two sub-regions were considerably far apart and were hence, considered as two distinct ROIs. We have also checked the consistency of these two sources at an individual subject level. In total, 18 out of the 19 participants showed activations in both the anterior and posterior right TPJ at 99^th^ percentile threshold while the remaining one participant exhibited the same upon further lowering the threshold to 95^th^ percentile.

### 3.4. Causal structure of the underlying sources

Source time series computed from 7 sources in dynamic stimulus and 5 sources in static stimulus processing conditions were subjected to Granger causality (GC) analysis to understand the directional influence between the sources. Out of all the significant functional connections (95% confidence interval) from a total of 42 possible connections during dynamic stimulus processing and 20 such connections during static stimulus processing, we chose to focus on the top 50 percentile connections that had a relatively high GC as illustrated in Figure 5.

**Figure 5.**
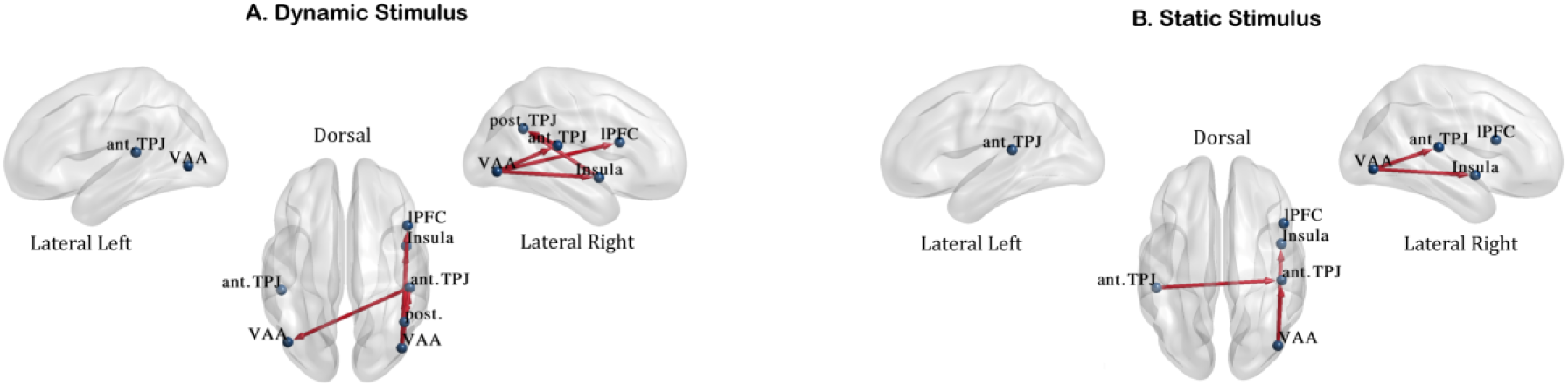
Effective Connectivity. The figure represents the directional influences between the localized sources responsible for processing a salient distractor for (A) Dynamic Stimulus and (B) Static Stimulus. The arrows point from the driver node towards the effector node. The causalities were determined using a time-varying Granger Causality approach on the source reconstructed time-series data and the figure illustrates the top 50% of all the significant causations for each stimulus condition.

Granger causality revealed the following causal influences:

Right Visual Area ➔ Right Insula, Right Visual Area ➔ Right anterior TPJ, Right Visual Area ➔ Right lateral PFC, Right Insula ➔ Right posterior TPJ, Right anterior TPJ ➔ Right posterior TPJ, Right anterior TPJ ➔ Left Visual Area for dynamic viewing and the following ones: Right Visual Area ➔ Right Insula, Right Visual Area ➔ Right anterior TPJ, Left anterior TPJ ➔ Right anterior TPJ for static viewing conditions.

The brain networks were visualized using BrainNet Viewer (Xia, Wang, & He, 2013). Overall, there is clearly a right hemispheric dominance of network interactions during saliency processing in both static and dynamic tasks.

## 4. Discussion

We investigated the importance of context specificity while processing saliency and the extent to which it reflects in the alpha modulation and its underlying neural networks. Our findings show: 1) The alpha power enhancement corresponding to the trials with saliency had common alpha peak power (∼10 Hz) with a near identical power rise for both the static and dynamic stimulus conditions. No other frequency band showed power changes between trials with saliency (ST) and without saliency (WT). This indicates a task-independent role of alpha oscillations while processing salient distractors, that does not depend on the task condition *per se* but is rather saliency-specific. 2) The underlying sources of the alpha oscillations obtained through EEG source localization were the right insula, the right lateral pre-frontal cortex (lPFC), the left and the right anterior temporo-parietal junction (TPJ) which are regions of the Ventral Attention Network (VAN) and were common to both the task conditions. These results further reiterate our earlier inference that the neural patterns of alpha enhancement vis-à-vis saliency and the underlying VAN are completely agnostic to the task conditions. 3) Not just the underlying sources but the effective connectivity between the sources also showed a similar pattern in both the task conditions where the right insula and the right anterior TPJ were driven by the right visual association areas, elucidating a posterior to anterior flow of bottom-up signals.

One of the possible explanations for such similar patterns in neural activity during saliency processing in both task conditions can be top-down influences that bias the competition between target and distractor by modulating bottom-up signals in a way that override the sensory-driven task-specific inputs.

### Neural oscillations and behavioral correlates of saliency processing

Reaction times of ST were significantly higher than reaction times of WT in both the task conditions reflecting the processing costs involved in the reorienting of attention to the salient distractors. Meanwhile, the NT, which had no possible correct responses and yet required a goal directed attention, resulted in the highest reaction times, possibly reflecting the role of increased cognitive load/ task complexity. Hence, it is important to characterize whether the neural dynamics corresponding to salient stimulus is due to an attentional shift alone or attention combined with task complexity.

Power spectral density results showed significantly enhanced levels of alpha power (8-12 Hz) for ST as compared to WT. Interestingly, enhancement in alpha power was not seen for NT (even though they had the highest reaction times) compared to WT, thus, ruling out the possibility of alpha power increase stemming from task complexity/difficulty level. This suggests that the underlying cause for the alpha power enhancement in the ST was distractor suppression (to improve task performance) which is consistent with the role of alpha oscillations from a vast body of literature (Fries, Reynolds, Rorie, & Desimone, 2001; Fu et al., 2001; Jensen & Mazaheri, 2010; Klimesch, 2012; Foxe, Simpson, & Ahlfors, 1998; Worden, Foxe, Wang, & Simpson, 2000; Foxe, Simpson, Ahlfors, & Saron, 2005; Snyder & Foxe, 2010; Banerjee, Snyder, Molholm, & Foxe, 2011; Zumer, Scheeringa, Schoffelen, Norris, & Jensen, 2014; Liu, Bengson, Huang, Mangun, & Ding, 2014; Feng, Störmer, Martinez, McDonald, & Hillyard, 2017). Also, a number of studies using attentional (spatial) cueing paradigms showed an increased amplitude in alpha band power for the to-be-ignored as compared with the to-be-attended location consistent with behavioral performance (Capilla et al., 2014; Capotosto et al., 2009; Feng et al., 2017; Foster et al., 2017; Frey et al., 2014; Ikkai et al., 2016; Kelly et al., 2006; Rihs et al., 2007; Schneider et al., 2019; Thut et al., 2006; Voytek et al., 2017; Worden et al., 2000). We not only replicated these findings but also extended the results for a task with spatio-temporal complexity as well.

### Underlying cortical sources of saliency processing across task conditions

The topoplots at peak alpha power (∼10 Hz) indicated the presence of a possible overlap in the right centro-parietal and parieto-occipital regions of cortex for processing saliency related information and guided us to further delve into the source space. Source reconstruction results revealed that most of the sources were common in both task conditions again indicating task-independent saliency processing. One might argue here that the relative increase of alpha power calculated using alpha modulation index might have negated task-specific information as well leaving behind stimulus properties which could only be attributed to the salient distractors. In this context, we would like to emphasize on the point that our dynamic and static task conditions consisted of salient distractors which too were dynamic and static, respectively which reinforces our claim for the involvement of common neural correlates in saliency processing across different stimulus conditions.

We observed that the right insula, the left and right anterior temporo-parietal junction (TPJ), the right lateral prefrontal cortex and the visual association areas showed source activations underlying alpha enhancement in both static and dynamic stimulus conditions. The aforementioned regions have been extensively shown to be involved in the processing of salient stimuli in previous fMRI studies and are also a part of the Ventral Attention Network (VAN) which is mostly right lateralized (Schuwerk, Schurz, Müller, Rupprecht, & Sommer, 2017; Eddy, 2016; Krall et al., 2015; Han & Marois, 2014). Though studies suggest that the VAN does not work in isolation but has a collaborative role with the Dorsal Attention Network (DAN) to process salient stimuli (Vossel et al., 2014), we chose to consider only VAN for our present study.

### The sub-regions of TPJ

Few studies have reported the involvement of both the left and the right TPJ while the right TPJ in particular, has been implicated in spatial reorienting (Chang et al., 2013) and in computing the behavioural relevance of salient signals (Geng and Mangun, 2011; Krall et al., 2015; Carter & Huettel, 2013; Decety & Lamm, 2007). This right-hemisphere asymmetry in TPJ activation is also seen in patients with hemineglect where there is a predominance of right rather than left TPJ lesions (Downar et al., 2001). The functional role of TPJ however, is highly debated and so is its location, due to an unusually high degree of inter-individual variability in its anatomical structure (Van Essen, 2005; Caspers et al., 2006) which makes the study of this region challenging. This may be due to the presence of different sub-regions of the TPJ involved in different functional roles (Bzdok et al., 2013). Hence, it is important to characterize the specific roles of these sub-regions, especially in the context of saliency processing.

An important observation from our source reconstruction results was the behavioral relevance specific posterior TPJ activation in the dynamic stimulus condition that is not seen in the static task condition. Earlier studies (Corbetta & Shulman, 2002; Corbetta et al., 2008) suggest that the right TPJ shows activation in response to salient distractors that are only “behaviorally relevant” (Huang, Tang, Sun, & Luo, 2018). A salient distractor can be considered to be behaviorally relevant if it shares features with the target of a task. The salient distractor in the dynamic task condition was hence behaviorally relevant to the task (differently sized/ colored dot amidst other dots) whereas in the static task condition it was not (complex naturalistic objects as opposed to a target which was a basic geometrical shape ‘+’). Using diffusion-weighted imaging tractrography–based parcellation, Mars and colleagues demarcated TPJ into 3 distinct sub regions: anterior TPJ (supra-marginal gyrus), posterior TPJ (angular gyrus) and the dorsal TPJ (middle part of the inferior parietal lobule) (Mars et al., 2012; Carter & Huettel, 2013; Geng & Vossel, 2013). Similar studies by Kubit and Jack (2013), identified these TPJ sub-regions to be associated with target detection, oddball identification and mentalization/ social cognition, respectively. The first two sub-regions are consistent with our source reconstruction results. Here, we note that such a crucial claim over the spatial anatomy of a sub-region just based on an EEG study comes with several limitations, nonetheless, underpinning the transient changes in neural dynamics during reorientation of attention associated with the functionally heterogeneous sub-regions of rTPJ, requires temporal precision which is difficult to achieve through other neuroimaging techniques like fMRI. Some recent evidences have shown that the detection of very focal activations for cortical sources may be possible using the source reconstruction algorithm eLORETA (Halder, Talwar, Jaiswal, & Banerjee, 2019). Furthermore, we have co-registered the EEG of individual participants with their respective brain MR images instead of directly warping the individual brain recordings to a standard template, in order to get a more accurate spatial estimate.

### Effective connectivity between the regions of VAN

In our study, identification of the almost common set of brain areas underlying alpha enhancement across dynamic and static stimuli warrants the understanding of the causal relationships among the candidate nodes. Causal influences from the visual association areas to the anterior (right) insula and anterior rTPJ in both the task conditions suggest that these two regions might be the key players in initiating the process of reorientation after receiving bottom-up signals from the visual areas to process a salient stimulus. The rTPJ is sensitive to stimuli that share target color (Hu et al., 2009, Natale et al., 2010, Serences et al., 2005) or category (Hampshire et al., 2007), and task cues that indicate a new target location (Shulman et al., 2009). An overall comparison of the causal architecture across static and dynamic stimulus conditions for an equal time-window suggests the active involvement of more number of sources to process saliency in the dynamic stimulus condition. This may be attributed to the spatio-temporal complexity of the task where the distractor changes its position at every instant on the time-scale. Previous studies have shown that the anterior insula is activated by the onset and offset of oddballs during attentional orienting/reorienting and is responsible for switching between large-scale networks to facilitate access to attention (Menon & Uddin, 2010) specifically in the context of perceptual decision making (Chand and Dhamala, 2016). Directional influence from the right insula to the posterior rTPJ in the dynamic stimulus condition might be indicative of driving such a switch in attention from VAN to DAN, required to get back to the goal-directed task. The possibilities and the consistency of these results however, need to be verified with several other experiments, particularly ones involving simultaneous EEG-fMRI. Functional connectivity patterns have shown that it is the anterior sub-region of TPJ that is mainly connected with the inferior frontal gyrus and the anterior insula constituting the VAN (Gillebert et al., 2013) and the lateral pre-frontal cortex (lPFC) plays a role in the integration of sensory information between the anterior insula and TPJ (Kubit & Jack, 2013). Granger causality analysis on dynamic viewing condition revealed that the posterior rTPJ is driven by the anterior rTPJ. Posterior TPJ has been shown to be associated with identification and evaluation based processes in oddballs (Kubit & Jack, 2013). The anterior TPJ on the other hand, is responsible for the reorientation of attention whenever faced with a salient distractor (Igelström et al., 2015; Rennig et al., 2015). If the salient distractor is behaviourally inconsequential, based on comparison of sensory information to internal representations (Krall et al., 2015), attention is oriented back to the task at hand. Based on our connectivity results, we propose that a behaviorally relevant salient distractor (as in the case of dynamic task) is further processed in the posterior rTPJ which evaluates the contents of the distractor and subsequently plays a gating role in driving the attentional requirements. Thus, the behaviourally relevant distractor is prioritized over a behaviorally irrelevant one at the stage of anterior TPJ such that the latter is not forwarded for further processing. On this view, a recent study has reported that the TPJ does not directly compute the relevance of a stimulus feature, but modulates its response to stimuli according to the top-down biasing of signals to control the engagement of attention to potentially distracting information that is behaviorally relevant to the task (Pedrazzini & Ptak, 2019).

Finally, as a prospective avenue for future research, we speculate that characterizing the various sub-regions of TPJ will probably be of critical importance to neuroscience in order to understand the subtle variations of attentional processing, both for spatial as well as spatio-temporally enriched stimuli. Although, various studies have explicitly mentioned about the sub-regions of TPJ in this regard, we would like to go a step forward and propose that the posterior TPJ receives top-down biasing signals from the anterior TPJ that help modulate our responses based on the relevance of a distractor to a putative task. Some previous studies have shown that abnormality in alpha oscillations correlates with attentional disorders. Atypical alpha asymmetry during covert attention has been observed in attention deficit hyperactivity disorder (ADHD) compared with neurotypical populations (Hale et al., 2009). Extending our study to other modalities like auditory and multisensory can establish communication in alpha frequencies via the VAN nodes as a marker of saliency processing which is agnostic to task complexity. Such a marker can prove to be of significant diagnostic use in clinical studies.

## Abbreviations

VAN: Ventral attention network
rTPJ: right temporo-parietal junction
NT: neutral trials
WT: without saliency trials
ST: saliency trials

## Acknowledgements

We thank Dr. Dipanjan Ray for his helpful comments on improving the readability of the manuscript, NBRC Core funds, and infrastructural support. PG was supported by Council of Science and Industrial Research (CSIR) fellowship (09/821(0044)/2017-EMR-I) from 03/08/2016, DR was supported by the Ramalingaswami fellowship (BT/RLF/Re-entry/07/2014) and DST-CSRI extramural grant (SR/CSRI/21/2016) and AB was supported by Ramalingaswami fellowship BT/RLF/Re-entry/31/2011) and Innovative Young Biotechnologist Award (IYBA), (BT/07/IYBA/2013).

